# Linking genomic signatures of selection to expression variation and direct evidence of local adaptation

**DOI:** 10.1101/2020.08.22.262394

**Authors:** Nicholas Price, Jack L. Mullen, Junjiang Lin, Christina Boucher, John K. McKay

**Affiliations:** Department of Bioagricultural Sciences & Pest Management, Colorado State University, Fort Collins, CO 80523, USA; Department of Computer and Information Science and Engineering, University of Florida, Gainesville, FL 32611, USA

## Abstract

Understanding how genomic and expression variation is linked to adaptation of plants to local environments is fundamental to the fields of evolutionary biology and species conservation. Using locally adapted *Arabidopsis thaliana* Italy and Sweden populations, we examine how variation in gene expression under control and cold acclimation conditions, is linked to allele frequency differentiation (AFD); linkage disequilibrium (LD); selective constraint at nonsynonymous sites; and genetic-tradeoff quantitative trait loci (GT-QTL). Our results indicate that contrary to genes showing a main effect in environment (E), expression genotype by environment interactions (GxE) show significantly higher AFD along cis-regulatory and nonsynonymous sites than the neutral expectation; and interestingly, highly differentiated GxE genes show higher expression and inter-species selective constraint than the rest of the genes. When examining the association between genomic signatures of selection along GxE/E genes and GT-QTL, we find that GxE genes showing a high AFD and LD, display a significant and much higher enrichment along GT-QTL than the genome-wide/E set of genes. Nonetheless, E genes show a higher enrichment than the genome-wide control. In summary, our results suggest, that these highly expressed and selectively constrained GxE genes, may have been part of a cold-responsive regulon of E genes that experienced recent selection when migrating to new environments. Candidate GxE genes underlying GT-QTL reveal interesting biological processes that may underlie local adaptation to temperature, including flowering time, light-dependent cold acclimation, freezing tolerance, and response to hypoxia. Finally, we find no evidence linking lower expression of the CBF-dependent freezing tolerance pathway to genetic-tradeoffs and adaptation to warmer climates.

## Introduction

Populations may vary in genotype, phenotype, and fitness across geographical regions that differ in abiotic variables of the environment. Such variation maybe generated after populations become adapted to local climates, where local genotypes have higher fitness than foreign genotypes at home (Kawecki and Ebert 2004; Hereford 2009; Des Marais, et al. 2013). Abiotic stress responsive gene expression (i.e., gene expression plasticity) may play a pivotal role in local adaptation, since studies have linked it to increases in stress tolerance (Rockman, et al. 2003; López-Maury, et al. 2008; Thomashow 2010; Brown, et al. 2017), and the adjustment of an organism’s life-cycle to favorable environmental conditions (Seo, et al. 2009; Chiang, et al. 2011). Understanding the link between genetic, expression, and fitness variation under different abiotic environments is central to the fields of evolutionary biology (Hoban, et al. 2016), conservation genomics (Razgour, et al. 2019), and plant breeding (Henry and Nevo 2014).

Gene expression responses under different environments, are often conserved between genotypes from different populations (Hannah, et al. 2006; Des Marais, et al. 2012), in which case, they will exhibit a main effect in environment (“E”) when plotted in a norm of reaction plot (Baye, et al. 2011). On the contrary, they may exhibit genotype-by-environment interactions (“GxE”), in which case the ranks of genotypes (G) change or switch from one environment to another (Baye, et al. 2011). E-genes may underlie adaptations to common environmental changes, while GxE-genes may underlie adaptation to aspects of the environment that significantly differ between locations within a specie’s range. For example, across the native range of a plant, all populations may face a slight deviation in temperature during winter/summer which engages a similar response among a set of genes of essential genes (i.e., E responses). On the other hand, parts of the native range may experience much harsher winters/summers than on average, thereby causing divergent selection between genotypes and the formation of GxE responses. If expression GxE interactions reflect fitness GxE interactions, then they may represent a important mechanisms of adaptation (López-Maury, et al. 2008; Franssen, et al. 2011; Morris, et al. 2014; Lovell, et al. 2016).

Genetic variation linked to expression and fitness GxE interactions can exhibit: (a) genetic tradeoffs, where the derived genotype is advantageous in one environment but deleterious in the other; and (b) conditional neutrality where the derived genotype is advantageous (conditionally advantageous) or deleterious (conditionally deleterious) in one environment and neutral in the other (Anderson, et al. 2011; Mee and Yeaman 2019). Despite the presence of fitness GxE interactions, instances of conditionally deleterious or more correctly non-locally maladaptive mutations, do not represent instances of ‘adaptation’ (Mee and Yeaman 2019). To identity candidate genetic variation underlying local adaptation at the single nucleotide level, the two of the main approaches used are: (a) identifying single nucleotide polymorphism (SNP) that show significantly higher allele frequency differentiation between populations than expected under neutral models of evolution (Beaumont and Balding 2004; Foll, et al. 2014; de Villemereuil and Gaggiotti 2015) and (b) identifying alleles showing significant associations to environment while accounting for population/geographic structure (Lasky, et al. 2012; Zhou and Stephens 2012; Gunther and Coop 2013; Luu, et al. 2017; Caye, et al. 2019). Loci underlying genetic-tradeoffs are expected to exhibit significantly stronger population genomic evidence of local adaptation than the genome average and conditionally advantageous loci (Tiffin and Ross-Ibarra 2014; Yoder and Tiffin 2017; Mee and Yeaman 2019).

Some of the main difficulties in identifying SNPs underlying local adaptation is disentangling adaptive, from neutral or slightly deleterious variation generated by background/relaxed selection, and genetic drift (Zhen and Ungerer 2008b; Hoban, et al. 2016; Matthey-Doret and Whitlock 2019). Simulation studies comparing various methods used to identify genetic variation underlying local adaptation while accounting for the effects of population structure have shown that the power of each method can significantly change depending on the underlying evolutionary scenario, in addition to other factors (De Mita, et al. 2013; de Villemereuil, et al. 2014; Lotterhos and Whitlock 2015; Yoder and Tiffin 2017). Furthermore, in examining the link between GWA/population-genomic methods and empirical evidence of local adaptation, the strength of this link changed depending on the method(s) used in studies (Fournier-Level, et al. 2011; Lasky, et al. 2014; Yoder, et al. 2014; Exposito-Alonso, et al. 2018; Price, et al. 2018; Price, et al. 2020).

Despite these hurdles, there have been many studies examining the genetic basis of local adaptation (Savolainen, et al. 2013; Hoban, et al. 2016), but only a few linking genome-wide expression variation, sequence variation, and fitness variation across selective gradients (Kelly 2019). Among the few, a study by Lasky, et al. (2014) examined the link between genomic signatures of local adaptation, patterns of expression, and fitness variation in Arabidopsis. The main result of the study was that expression GxE genes showed a higher enrichment of climate-correlated SNPs than genes showing a main effect in environment (E); suggesting a role of expression GxE interactions in local adaptation (Lasky, et al. 2014). Nonetheless, the enrichment of fitness associations along GxE genes was not significant despite being higher than E genes. This discordance could be the result of differences in purifying selection between E and GxE genes leading to an enrichment of slightly deleterious climate-associated SNPs in the latter set (Mee and Yeaman 2019).

Arabidopsis wild populations offer a valuable resource to re-examine the interplay between genetic, expression, and fitness variation across climatic conditions. The native range of Arabidopsis includes parts of Northern and Southern Europe that experience significantly different climatic conditions. For example, populations in North Sweden, experience average soil temperatures below freezing for about a 1/3 of the year, while in North-Central Italy such temperatures are rarely recorded (Oakley, et al. 2014). Reciprocal transplant experiments have showed strong adaptive differentiation between these populations and evidence of genetic-tradeoffs (Ågren and Schemske 2012; Ågren, et al. 2013). Among the traits suggested to underlie these genetic-tradeoffs, is freezing tolerance (Oakley, et al. 2014); and more specifically freezing tolerance variation associated with the CBF pathway (Thomashow 2010; Park, et al. 2015; Park, et al. 2018). Studies have suggested that the lower freezing tolerance and expression of this pathway (Cook, et al. 2004; Hannah, et al. 2006; McKhann, et al. 2008; Gehan, et al. 2015) across Arabidopsis populations in warm climates (e.g., Italy), is an adaptive response that is deleterious in cold climates (e.g., Sweden) (Oakley, et al. 2014). This adaptive response has been linked to non-functionalization of the CBF-pathway (Oakley, et al. 2014; Gehan, et al. 2015; Monroe, et al. 2016). Nonetheless this hypothesis has been disputed by studies showing that this nonfunctionalization and decrease in freezing tolerance is due to relaxed selection in warmer climates (Zhen and Ungerer 2008a, 2008b; Zhen, et al. 2011).

To re-examine the link between genetic, expression, and fitness variation of Arabidopsis populations in different climates, and the role of the CBF-pathway in local adaptation to temperature, the current study examines the following data: (a) re-sequenced genomes of locally adapted (Ågren and Schemske 2012) South Italy and North Sweden (Price, et al. 2020); (b) expression of Italy and Sweden genotypes under control and cold-acclimation conditions (Gehan, et al. 2015); and (c) quantitative trait loci (QTL) explaining fitness variation of Italy and Sweden recombinant inbred lines grown in a series of reciprocal transplant experiments (Ågren, et al. 2013). More specifically, we examine the link between allele frequency differentiation (AFD) and linkage disequilibrium (LD) at cis-regulatory (sites found 1kb upstream from the transcriptional start site) and nonsynonymous sites, to patterns of expression (E and GxE) and genetic-tradeoff QTL, while taking into account the effects of selective constraint (or purifying selection) at nonsynonymous sites. Among population genomic signatures of local adaptation we chose AFD and LD, since these were previously found to be enriched along fitness QTL (Price, et al. 2020).

## Materials and Methods

### Extraction of RNA under cold conditions and sequencing

The Arabidopsis SW and IT accessions were collected from their native habitats in Sweden and Italy, respectively (Ågren and Schemske 2012). Plants were grown at 22°C on soil under a 12 h photoperiod for 18–26 days (control), or at 4°C under a 12 h photoperiod for 1 or 2 weeks (cold treatment). Rosette tissue was collected from plants exposed to low temperature (4°C) for 0, 1, and 2 weeks. Total RNA was isolated for each experimental replicate (three replicates). Nine replicates were collected for each accession, for a total of 18 biological replicates. Further details can be found in the study by Gehan et al. (2015).

RNA was submitted for RNAseq library prep and 100bp single-end RNAseq analysis to Michigan State University’s Research Technology Support Facility (RTSF). Sample preparation was performed by MSU RTSF with standard protocols of the mRNA-Seq Sample Preparation Kit (Illumina). Sequencing was performed on an Illumina Genome Analyzer II (Illumina). Three samples were multiplexed in a lane for a total of 6 lanes. After quality trimming, RNAseq resulted in single-end reads ~75 bp in length with an average of 45,257,092 reads passing the Illumina purity filters for each sample. To map reads to the *Arabidopsis thaliana* genome we used Tophat (Trapnell, et al. 2009) and we estimated transcript abundance using Cufflinks (Trapnell, et al. 2010).

### Identifying differentially expressed genes between Italy and Sweden accessions

To identify genes showing a main effect in environment (or condition) (E) and genotype by environment interactions (GxE) we used the we used the package DESeq2 (Love *et al*. 2014) and focused on expression after one week of cold. More specifically, using the function “DESeqDataSetFromMatrix” and a design to identify genotype by environment interactions (“genotype+condition+genotype*condition”) we identified GxE genes that showed an adjusted p-value (“padj”) of <0.01. Thereafter, using the “contrast” argument we extracted genes that showed a main effect in environment (E) using a padj <0.01. To ensure no main ffect in genotype among E genes we removed any genes showing a main effect in genotype (G) using a p-value of 0.05.

### Comparing mean expression and selective constraint across E and GxE genes

To compare average expression of Italy and Sweden plants across E and GxE genes we first estimated the average “Fragments Per Kilobase of exon model per Million mapped fragments” 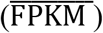 of the three samples under each pair of conditions (“control”, “cold”). Using the average expression of each gene under control and cold conditions, we estimated the mean expression and 95% CI’s of all genes in each category (i.e., E and GxE genes). To estimate 95% CI’s we used10,000 bootstrap samples. In addition to average expression of genes across each condition, we also estimate average difference in expression between conditions 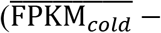 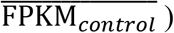. Selective constraint/ purifying selection at nonsynonymous sites was examined using the ratio of nonsynonymous to synonymous rates of substitution (dN/dS). dN/dS ratios were downloaded from EnsemblPlants (Howe, et al. 2020) Biomart (Kinsella, et al. 2011), using *Arabidopsis thaliana* and *Arabidopsis halleri* orthologs. dN/dS ratios above 1 were ignored.

### Population genomic signals of selection

As population genomic signatures of local adaptation we used a combination absolute allele frequency differentiation 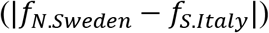 and linkage disequilibrium (LD) between a SNP and its neighboring SNPs with a 20kb window. LD was measured using the package ‘PLINK’ (Purcell, et al. 2007) and it was estimated as the mean square coefficient of correlation 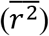. AFD and LD were estimated in a previous study (Price, et al. 2020).

### Defining cis-regulatory and nonsynonymous variation

Cis-regulatory sites of genes were defined using a maximum length of 1 kb from the transcriptional start site unless there was overlap with the transcribed region of another gene in which case the promoter region was shorter. For sites that were associated to two genes, were assigned to the nearest gene. To call nonsynonymous variation among Italy and Sweden accessions we used bi-allelic sites, a publicly available python script (callSynNonSyn.py; archived at https://github.com/kern-lab/), and gene models downloaded from the TAIR database (TAR10 genome release) (Berardini, et al. 2015).

### Circular permutation tests to examine evidence of local adaptation across groups of genes

To examine whether the proportion of (E/GxE) genes with cis-regulatory/nonsynonymous SNPs showing evidence of local adaptation (estimated using AFD and/or LD) is significantly higher than expected by chance we used a circular permutation test. This test has been previously explained in detail (Price, et al. 2020); but in brief, AFD’s and/or LD’s are shifted across the genome (not randomly shuffled) and according to certain criteria (e.g., AFD>0.60) we estimate the proportion of genes with high AFD and/or LD cis-regulatory/nonsynonymous SNPs. This was repeated a thousand times and the resulting permutation distribution is compared to the observed proportion/number of genes with high AFD.

### Assembling CBF-regulon genes and flowering time estimates

Genes predicted to be regulated by the three CBF transcription factors (CBF’s 1-3) were retrieved from Park, et al. (2018) resulting in a set of 476 genes. Estimates of flowering time for 835 Eurasian *A. thaliana* accessions were downloaded from the study by Alonso-Blanco et. al (2016) (1001 Genomes Consortium 2016).

## Results

### Sweden but not Italy plants show significant upregulation of GxE genes under cold

To examine gene expression in Italy and Sweden plants under control (22 °C) and cold conditions (4 °C) we used DESeq2 (Love, et al. 2014). Using an FDR of <0.01, we identified 392 that showed genotype by environment interactions (“GxE”), and 2, 883 that showed a main effect in environment (“E”) after removing genes that showed a main effect in genotype at a p-value <0.05. To compare mean expression of these genes between Italy and Sweden plants we estimated mean Fragments Per Kilobase Of Exon Per Million Fragments Mapped 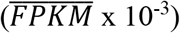 under control and cold conditions and determined the 95% CI’s using a bootstrap approach (Figures 1a & 1b). As expected, mean expression of “E” genes under control and cold conditions was identical in Italy and Sweden plants (Figure 1a). On the other hand, mean expression of “GxE” genes was significantly different between Italy and Sweden plants under cold conditions (Figure 1b); with Sweden plants showing a significant upregulation of genes when compared to control conditions 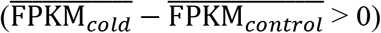 (Figure 1c). On the contrary, Italy plants showed a decrease in expression under cold conditions when compared to control conditions 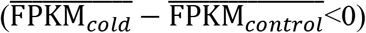 (Figure 1c).

**Figure 1.**
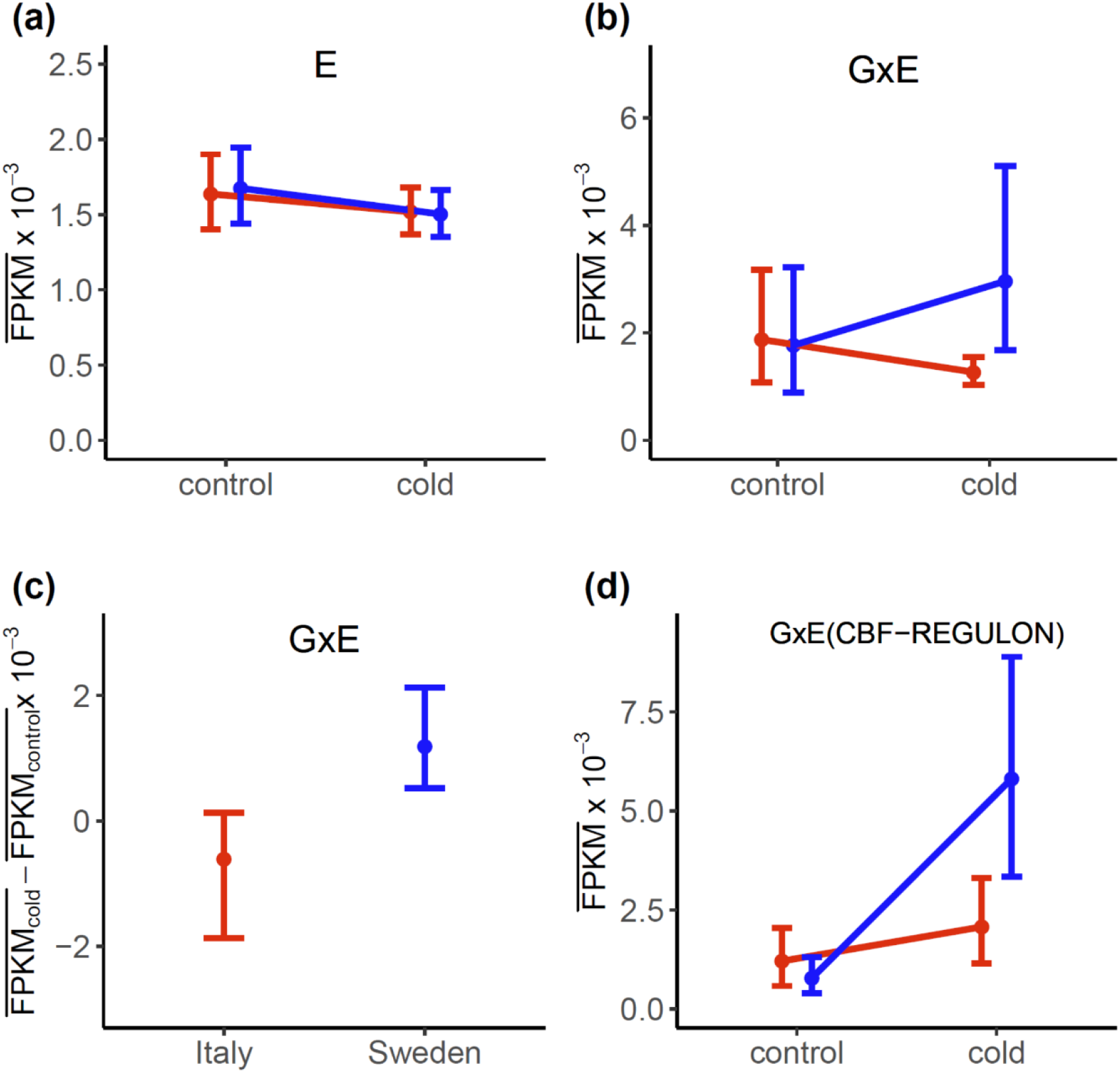
(a-b) Mean expression 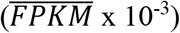 of E and GxE genes under control and cold conditions (FPKM: Fragments Per Kilobase Of Exon Per Million Fragments Mapped). Shown are the means and 95% confidence intervals estimated using a bootstrap approach. Mean expression of Sweden plants under cold conditions was significantly higher than in Italy plants (c) In comparison to control conditions 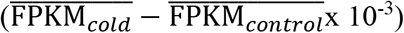 Sweden plants showed a net upregulation (≫0) under cold, while Italy plants showed a net downregulation (≪0) (d) Despite the opposite trends (c), mean expression of CBF-regulon GxE genes in Italy and Sweden plants, was higher under cold than control conditions; with Sweden plants showing the largest increase (>2 times higher).

To examine whether this decrease was also present among known freezing tolerance genes we examined mean expression of 53 CBF-regulon genes that showed GxE interactions. As shown in Figure 1d, expression of GxE CBF-regulon genes was significantly higher under cold than control conditions, in both Italy and Sweden plants. The increase in Sweden plants, however, was twice as high than in Italy plants. Among the set of 476 CBF-regulon genes (Park, et al. 2018), we identified 23 “E” genes and 53 “GxE” genes. In comparison to the total number of “E” and “GxE” genes (E: 2883, GxE: 392) CBF-regulon genes showed a significant enrichment (p-value<0.01) in GxE interactions according to single tail fisher’s test (“fisher.test” implemented in R).

### GxE but not E genes show significant allele frequency differentiation at cis-regulatory and nonsynonymous sites

Loci underlying local adaptation are expected to show significant allele frequency (Tiffin and Ross-Ibarra 2014) between populations. To examine such evidence along E and GxE genes, we looked at allele frequency differentiation at cis-regulatory and nonsynonymous SNPs. As a measure of allele frequency differentiation we used absolute allele frequency difference of the non-reference allele between Italy and Sweden populations 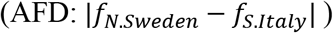 (Berner 2019; Price, et al. 2020); which showed a strong correlation (R^2^: 0.96, Figure S1) with a two population, two allele, F_ST_ measure (Bhatia, et al. 2013).

Along candidate cis-regulatory sites, the observed proportion of E genes (Figure 2a) with Cis-regulatory SNPs showing different levels of AFD’s (AFD<0.40, AFD>0.60, 0.80) did not significantly differ than the expected proportion (Figure 2a). On the other hand, the proportion of GxE genes with cis-reg. SNPs showing low AFDs (<0.40) was significantly lower than the expectation, while the proportion associated with high AFD’s (>0.60, 0.80) was significantly higher (Figure 2a). The difference in AFD’s along cis-regulatory SNPs can also be seen when comparing the histograms of E and GxE genes. As shown in Figure 2b, GxE genes show a higher proportion of cis-reg. SNPs with high AFD’s (>0.60) than E genes. The two histograms were found to be significantly different according to a X^2^ test (X-squared = 25.177, df = 9, p-value = 0.0028).

**Figure 2.**
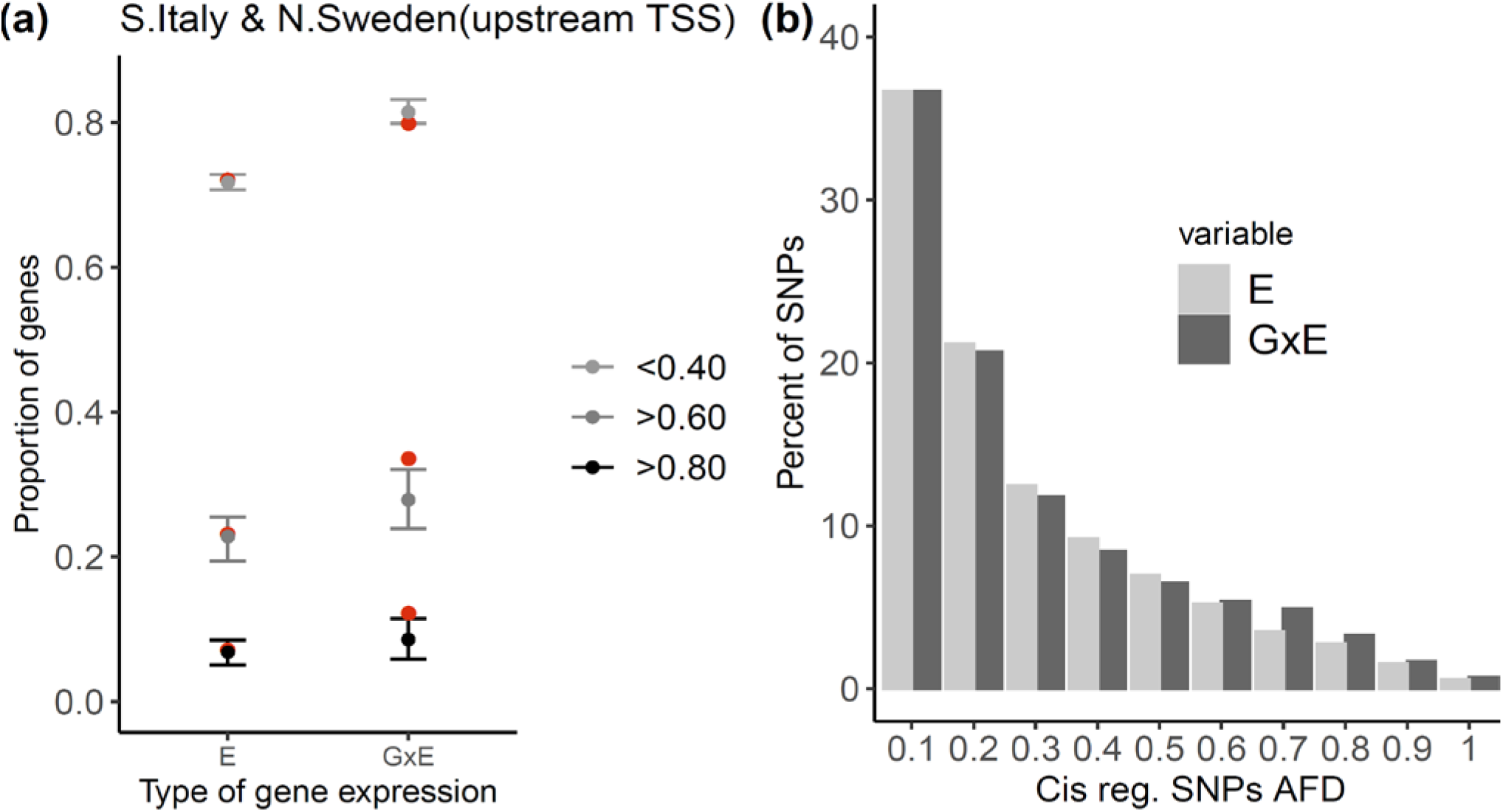
(a) The proportion of GxE genes showing high allele frequency divergence (AFD) at cis-regulatory sites was significantly higher than the expectation derived using circular permutations of genome wide SNPs. E genes on the other hand, showed no significant genetic differentiation along these sites. (b) The significantly higher differentiation is also observed when comparing the AFD distributions along cis-regulatory sites of E and GxE genes (X-squared = 25.177, df = 9, p-value = 0.0028). GxE genes show a higher proportion of cis-regulatory SNPs with an AFD>0.60.

To test whether nonsynonymous variation across “E” and/or “GxE” genes showed evidence of local adaptation we compared AFD distributions of nonsynonymous and synonymous variation. As depicted in Figure 3a, the distribution of AFD’s at nonsynonymous and synonymous sites of “E” genes is not significantly different according to a X^2^ test (X^2^: 11.0, df:9, p-value: 0.28). Contrary to “E” genes, the distribution of AFD’s at nonsynonymous sites of “GxE” genes was significantly different (X^2^: 18.5, df: 9, p-value: 0.03) than synonymous sites (Figure 3b). More specifically, nonsynonymous sites across “GxE” genes showed an enrichment in low AFD’s and high AFD’s (Figure 3b) which can be caused by recent local adaptation and/or purifying selection (Nielsen 2005).

**Figure 3.**
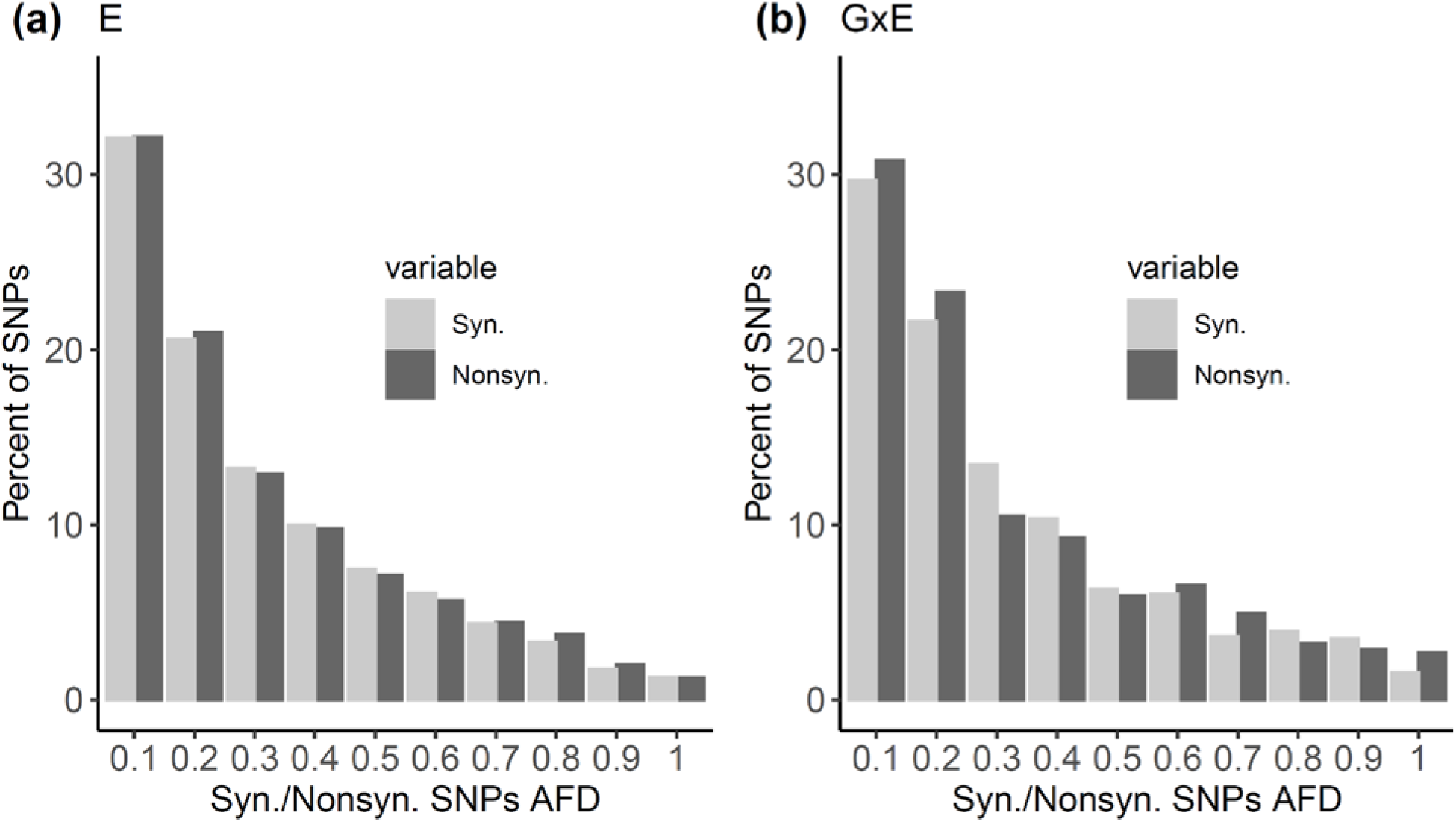
(a) Allele frequency divergence (AFD) between synonymous and nonsynonymous sites of E genes did not significantly differ according to a X2 test (X-squared = 11.002, df = 9, p-value = 0.28). (b) When comparing AFD between synonymous and nonsynonymous sites of GxE genes identifies a statistical significance (X: 18.546, df = 9, p-value = 0.029). The proportion of nonsynonymous SNPs with AFD’s <0.2 and >0.6 was higher than synonymous sites. Such deviations could be caused by both recent selection and purifying selection.

### Contrasting patterns of expression and purifying selection between GxE and E genes exhibiting low and high AFD

Variation in allele frequency differentiation between nonsynonymous sites of genes can result from differences in purifying selection, in addition to local adaptation. Furthermore, these differences could be associated with variation in gene expression among genes. To examine the link between AFD, purifying selection, and expression we first split E and GxE genes into ones that showed an AFD>0.60 at least one cis-regulatory/nonsynonymous site (AFDhigh) and ones that did not (AFDlow); furthermore, we narrowed down the sets of genes to those that we had estimates of dN/dS (dN: rate of nonsynonymous substitutions per site; dS: rate of synonymous substitutions per site); a measure of purifying selection at nonsynonymous sites.

E genes with high AFD SNPs showed a mean 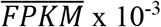 of ≈1 across all conditions and ecotypes, while E genes with low AFD showed an 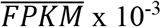 significantly higher than 1 (Figures 4a-4b). The opposite trend was seen across GxE genes. GxE genes with low AFD SNPs showed significantly lower expression 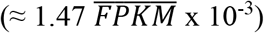 across the two conditions and ecotypes, than GxE genes with high AFD SNPs 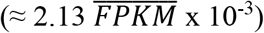 (Figures 4c-4d). In association with the levels of expression we observed a change in patterns of selective constraint/purifying selection along protein coding genes. As shown in Figure 5, E genes that contained high AFD SNPs, and were expressed at lower levels than AFDlow E genes, also showed a higher dN/dS, which translates to lower levels of purifying selection. On the other hand, AFDhigh GxE genes showed higher expression than AFDlow GxE genes, and a lower dN/dS (Figure 5).

**Figure 4.**
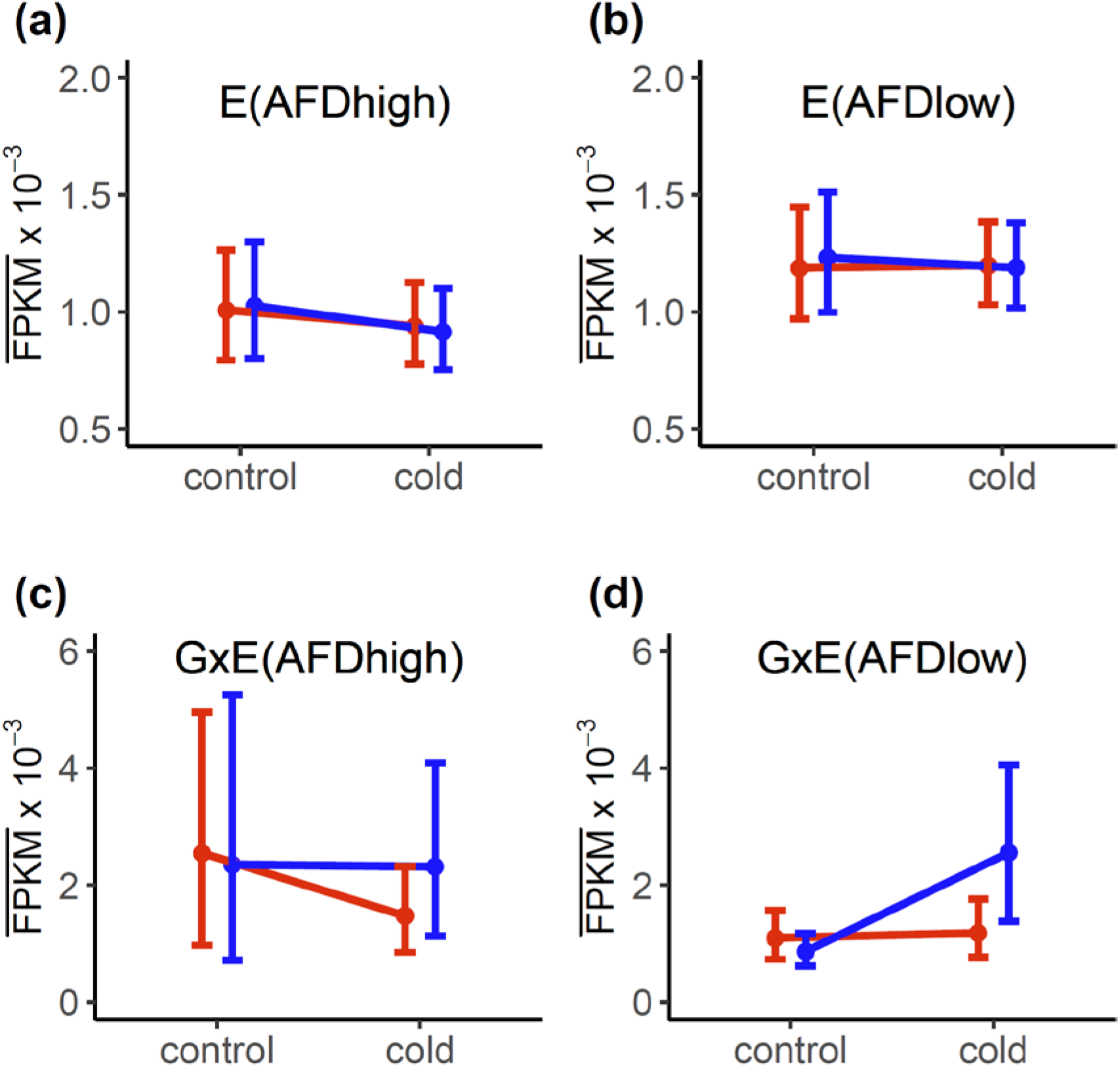
(a-b) E genes with at least one nonsynonymous/cis-regulatory SNP showing an AFD>0.6 (AFDhigh) were expressed at significantly lower levels when compared to the rest of the E genes with SNPs at cis-regulatory/nonsynonymous sites (AFDlow). Mean expression 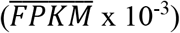 of AFDhigh E genes across ecotypes and conditions was ≈1, while that of AFDlow E genes was ≫1. (c-d) On the other hand, AFDhigh GxE genes showed a higher mean expression 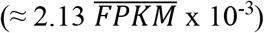 across ecotypes and conditions than AFDlow GxE genes 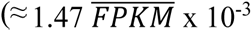.

**Figure 5.**
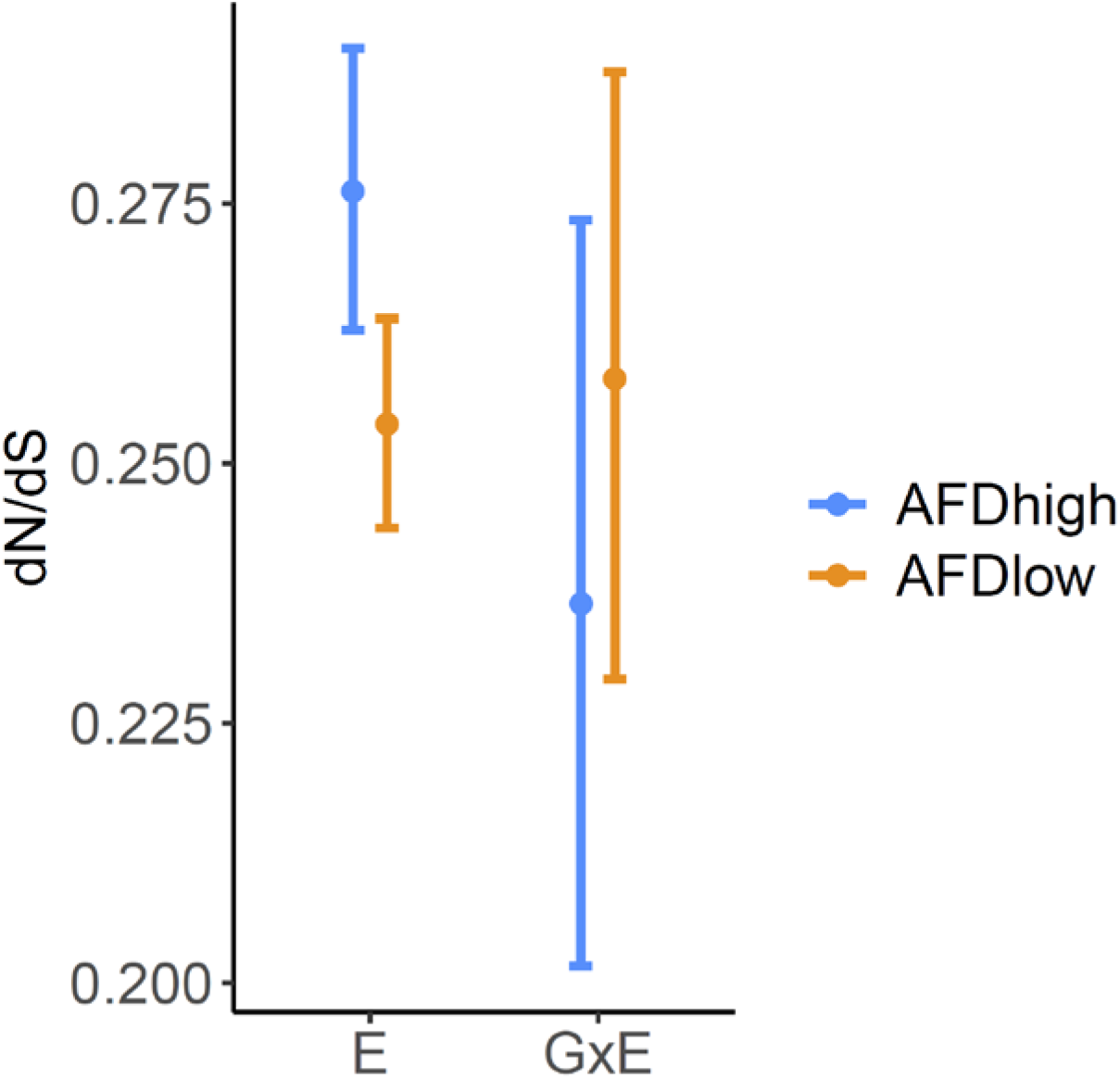
Patterns of purifying selection differed significantly between GxE and E genes that showed different levels of allele frequency divergence (AFD) at nonsynonymous sites/cis-regulatory sites. E genes with high AFD SNPs (AFDhigh) showed significantly lower purifying selection (higher dN/dS) at nonsynonymous sites, than E genes with low AFD SNPs (AFDlow) and GxE genes with high AFD SNPs (AFDhigh). GxE genes with high AFD SNPs also showed a lower dN/dS than GxE genes with low AFD SNPs. These patterns of dN/dS seem to correlate with the level of expression associated with these genes (Figure 4). The ration dN/dS was estimated using pairs of *A. thaliana* and *A. halleri* orthologous genes.

### Significant evidence linking GxE genes to recent selection and genetic tradeoffs

So far, we have showed that GxE genes showed significantly higher genetic differentiation than E genes at both cis-regulatory and nonsynonymous sites, in addition to higher levels of expression and purifying selection (Figures 2-5). To examine the association of E and GxE genes to fitness variation of Italy and Sweden genotypes at native sites, we used a set of 20 fitness QTL that were previously assembled into six genetic trade-off QTL that spanned ~18.6 Mbs of the genome (Ågren, et al. 2013). More specifically, we examined the distribution of genes showing high allele frequency differentiation (AFD) and linkage disequilibrium (LD) along genetic tradeoff QTL.

As shown in Table 1, in most instances the proportion of E genes withing genetic-tradeoff QTL was significantly higher than the genome-wide proportion; even when not filtering for high AFD and LD. On the other hand, GxE genes did not show any significant enrichment when not filtering for a high LD (Table 1). On the other hand, when we filtered for a high AFD (>0.60) and LD (>019, 0.32) (0.32 represents the 99^th^ percentile of the genome wide distribution of LD) the proportion of GxE genes along genetic-tradeoff QTL was significantly higher than the genome-wide proportion and the proportion of E genes satisfying these criteria (Table 1). In addition to examining the proportions along the six genetic tradeoff QTL, we also examined the proportions of these genes within 100kb of fitness QTL peaks (Table S1). The only significant enrichment was observed when examining the proportion of GxE genes with a high AFD (>0.60) and LD (>0.32) (Table S1). This proportion was significantly higher than the genome-wide set of genes, and E genes (Table S1).

**TABLE 1.** Comparing the proportion E and GxE genes exhibiting different signatures of local adaptation and selection, and found within six genetic tradeoff (GT) QTL peaks explaining fitness variation between Italy and Sweden populations (Ågren, et al. 2013). The table splits a set of genome-wide, E, and GxE genes according to their location along GT QTL and signatures of selection (AFD: Allele frequency divergence & LD: Linkage disequilibrium). The odd ratios depicted are derived by comparing the proportion of E/GxE genes to the genome-wide set of genes. A significantly higher proportion relative to the genome-wide set is indicated by a *, and significantly higher proportion when comparing E and GxE genes is indicated by a †. Comparison of proportions was done using a fisher’s one-tail test (p-val<0.05) implemented in R (“fisher.test”).

|  | NON-QTL |  |  | GT-QTL |  |  | E<br>(Odds-<br>ratio) | GxE<br>(Odds<br>ratio) |
| --- | --- | --- | --- | --- | --- | --- | --- | --- |
|  | Genome-wide | E | GxE | Genome-wide | E | GxE |  |  |
| No filtering | 24253 | 2166 | 305 | 5725 | 624 | 74 | 1.22*† | 1.03 |
| AFD$>0.60$ | 9082 | 734 | 128 | 2343 | 246 | 31 | 1.30*† | 0.94 |
| AFD$>0.60$ &<br>LD$>0.19$ | 1209 | 105 | 11 | 419 | 49 | 11 | 1.35* | 2.88*† |
| AFD$>0.60$ &<br>LD$>0.32$ | 192 | 22 | 0 | 71 | 9 | 6 | 1.10 | Inf*† |

To identify potential candidates underlying genetic tradeoffs, we chose GxE genes with cis-regulatory/nonsynonymous SNPs with high AFD (>0.60) and LD (>0.19) (Table 2). As shown in Table 2, GxE genes within genetic tradeoff QTL showed twice the expression levels than GxE genes outside the QTL. Two of these genes (AT2G35050, *FLDH*) were previously identified as candidate genes (Price, et al. 2018; Price, et al. 2020) and four of the genes (AT2G35050, AT3G56408, AT4G33180, AT5G65860) have no known function. The rest of the genes have been associated with some very interesting biological processes, such as shade avoidance, light-dependent cold tolerance, drought and freezing tolerance, and response to hypoxia (Table 2). *COL7* which is located within genetic tradeoff QTL 1:3, is also located within a high confidence flowering time QTL (Chr1: 27.4-29.1 Mb) where the Sweden genotype showed significantly longer flowering time than the Italy genotype in Italy (Ågren, et al. 2017). As shown in the rooted *COL7* tree (Figure S2) Eurasian accessions with a similar sequence at the Swedish genotype flo wer significantly later, than accessions with sequences as the Italy genotype.

**TABLE 2.**
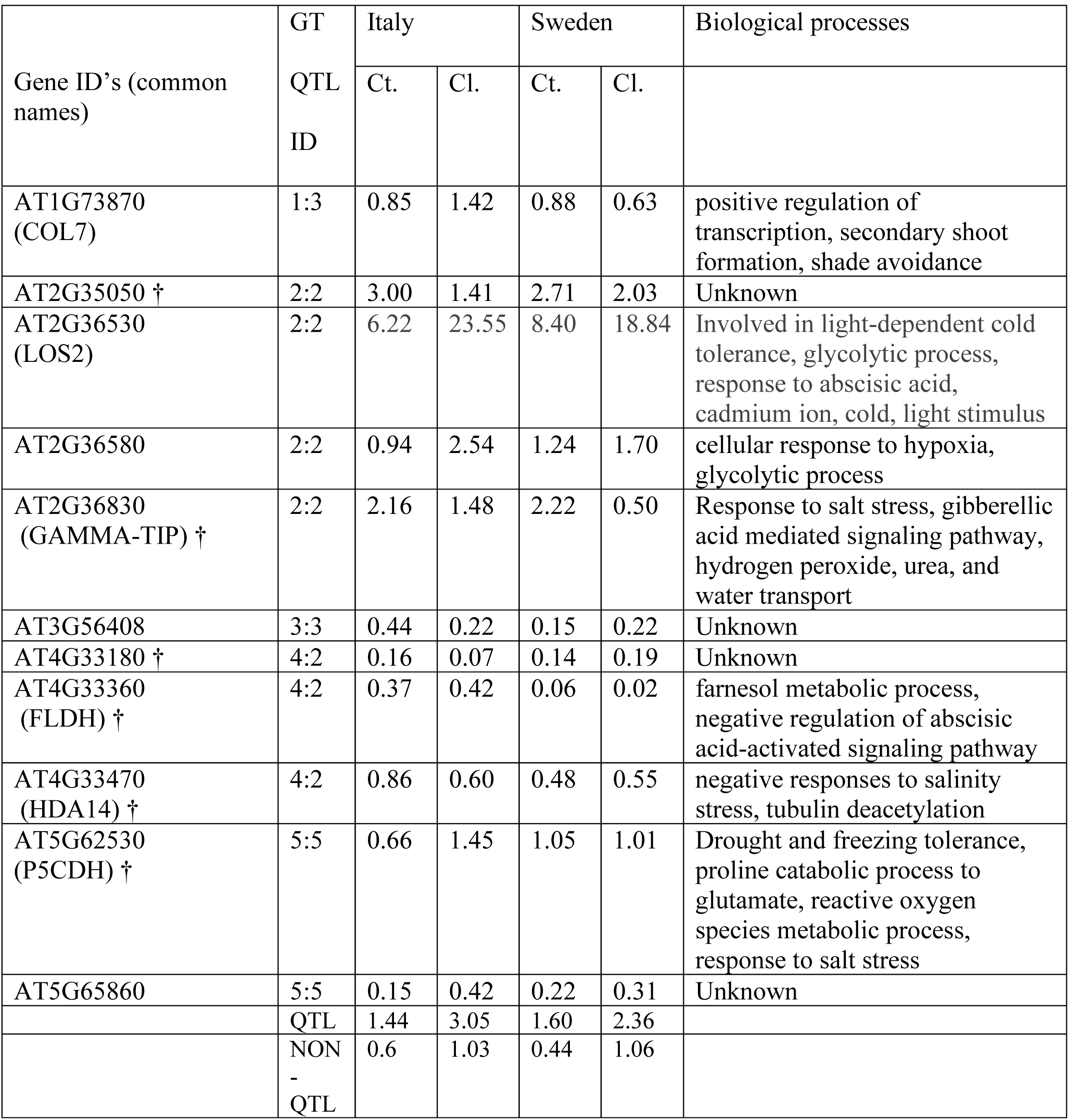
Genes showing significant genetic differentiation (AFD>0.60) and linkage disequilibrium (LD>0.19,0.32) along genetic tradeoff (GT) QTL (ID’s shown from Ågren, et al. 2013). Shown is also the mean expression 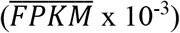 of these genes under control (Ct. – 22 °C) and cold acclimation conditions (Cl.-4 °C). † indicates genes with cis-reg./nonsyn. SNPs with an AFD>0.60 and LD>0.32). The Biological process were taiken from the TAIR database (Berardini, et al. 2015).

| Gene ID's (common names) | GT QTL ID | Italy |  | Sweden |  | Biological processes |
| --- | --- | --- | --- | --- | --- | --- |
|  |  | Ct. | Cl. | Ct. | Cl. |  |
| AT1G73870<br>(COL7) | 1:3 | 0.85 | 1.42 | 0.88 | 0.63 | positive regulation of transcription, secondary shoot formation, shade avoidance |
| AT2G35050 † | 2:2 | 3.00 | 1.41 | 2.71 | 2.03 | Unknown |
| AT2G36530<br>(LOS2) | 2:2 | 6.22 | 23.55 | 8.40 | 18.84 | Involved in light-dependent cold tolerance, glycolytic process, response to abscisic acid, cadmium ion, cold, light stimulus |
| AT2G36580 | 2:2 | 0.94 | 2.54 | 1.24 | 1.70 | cellular response to hypoxia, glycolytic process |
| AT2G36830<br>(GAMMA-TIP) † | 2:2 | 2.16 | 1.48 | 2.22 | 0.50 | Response to salt stress, gibberellic acid mediated signaling pathway, hydrogen peroxide, urea, and water transport |
| AT3G56408 | 3:3 | 0.44 | 0.22 | 0.15 | 0.22 | Unknown |
| AT4G33180 † | 4:2 | 0.16 | 0.07 | 0.14 | 0.19 | Unknown |
| AT4G33360<br>(FLDH) † | 4:2 | 0.37 | 0.42 | 0.06 | 0.02 | farnesol metabolic process, negative regulation of abscisic acid-activated signaling pathway |
| AT4G33470<br>(HDA14) † | 4:2 | 0.86 | 0.60 | 0.48 | 0.55 | negative responses to salinity stress, tubulin deacetylation |
| AT5G62530<br>(P5CDH) † | 5:5 | 0.66 | 1.45 | 1.05 | 1.01 | Drought and freezing tolerance, proline catabolic process to glutamate, reactive oxygen species metabolic process, response to salt stress |
| AT5G65860 | 5:5 | 0.15 | 0.42 | 0.22 | 0.31 | Unknown |
|  | QTL | 1.44 | 3.05 | 1.60 | 2.36 |  |
|  | NON<br>-<br>QTL | 0.6 | 1.03 | 0.44 | 1.06 |  |

### No evidence linking large decreases in mean expression of cold-induced and CBF pathway genes do adaptation in warm climates

Mean expression of high AFD and LD (AFD>0.60 & LD>0.19) GxE genes that showed significant enrichment along genetic tradeoff QLT did not show significant difference in mean expression (Table 1), as reflected in Figure 4c. The filtering for high LD maybe linked to instances of genetic tradeoffs but removes instances conditional neutrality where linkage disequilibrium is expected to be weaker but nonetheless higher than the neutral expectation. To test whether conditional neutrality is linked to the lower expression of some GxE genes in Italy (Figure 4c), we removed the genes in Table 2 from the set of high AFD GxE genes, and tested whether there was a significant difference in LD between 89 genes where Italy plants showed lower expression than Sweden plants under cold, and the rest of the 48 genes. Mean LD of these sets of genes was approximately the same (LD ~0.09), and not significantly different than the genome average (LD ~0.06), at cis-regulatory and nonsynonymous SNPs of LD<0.19. In addition to just choosing genes with a lower expression in Italy, we also looked at LD across genes where Italy plants showed lower or equal to half the expression of Sweden plants under cold. This resulted in 21 (out of the 89) genes, that also showed approximately the same LD (~0.09).

To further examine the possible role of the CBF pathway in causing genetic-tradeoffs we examined the proportion of CBF-regulon genes with high AFD, or high AFD and LD, cis-regulatory/nonsynonymous SNPs within genetic tradeoff QTL (Table S2). Relative to the genome-wide set of genes none of the categories examined showed a significant enrichment along the QTL (Table S2). In addition to CBF-regulon genes, we also examined whether the genomic region that included the three CBF transcription factors (CBF1-3), showed any genetic differentiation. Using a sliding-window approach we examined the proportion of high AFD (>0.60) SNPs along these genes (Figure S3). The proportion of high AFD SNPs across these genes was below the genome average (Figure S3). Under the assumption of genetic-tradeoffs we would expect these regions to show a significant increase in allele frequency differentiation (Tiffin and Ross-Ibarra 2014). Furthermore, we did not find any significant evidence for recent selection, since 19 cis-regulatory (no nonsynonymous SNPs) SNPs of the three CBF genes showed a very low mean LD (~0.05).

## Discussion

The current study re-examines the link between genome-wide sequence and expression variation to fitness variation of Arabidopsis populations showing significant evidence of local adaptation in their native environments (Ågren and Schemske 2012; Ågren, et al. 2013). The enrichment of genes showing a main effect in environment (E) along the low-resolution fitness QTL, in combination with the even higher enrichment of GxE genes showing significant genetic differentiation and linkage disequilibrium, suggest that plastic responses play an important role in adaptation. More specifically, E genes may represent regulon of genes that are necessary for facing common environmental challenges, while GxE genes represent instances of loci that underwent divergent evolution to adapt to extreme environmental differences. Furthermore, our results suggest that local adaptation occurs through highly expressed and selectively constraint genes. Finally, we find no significant evidence linking significantly lower expression of the CBF-pathway, to adaptation to warmer climates.

Local adaptation is expected to cause allele frequency differentiation (AFD) between populations; especially in the case of genetic tradeoffs (Tiffin and Ross-Ibarra 2014). Furthermore, if local adaptation is recent, loci should also exhibit high linkage disequilibrium (LD) (Nosil, et al. 2009). The importance of these signatures were shown in a previous Arabidopsis study (Price, et al. 2020), where we found that high AFD and LD SNPs were enriched along fitness QTL, and genes involved in life-history traits that are thought to play a major role in local adaptation; such as flowering time. One of these genes (*PIF3*) showed significant evidence of local adaptation in an evolutionary distant species of tree (*Populus balsamifera L.*) (Keller, et al. 2012). In the current study, we find that expression GxE genes with high AFD and LD SNPs are significantly enriched along individual fitness QTL and paired genetic tradeoff QTL, providing strong evidence for their possible role in local adaptation. Candidate GxE genes were tied to interesting biological processes such as: flowering time, light-dependent cold acclimation, freezing tolerance, and response to hypoxia.

Our final set of GxE genes with high AFD and LD did not show any significant differences in mean expression between Italy and Sweden (Table 2). On the other hand, high AFD GxE genes with lower LD (LD<0.19) showed large decreases in mean expression under cold in Italy plants (Figure 4C). These genes, however, did not show any enrichment along genetic-tradeoff QTL, and significant increases in LD. Some of the factors that may contribute to this observation, are that some of these expression interactions are neutral/nearly-neutral, or they involve adaptive mutations of small effect size (Yeaman 2015; Hoban, et al. 2016; Forester, et al. 2018; Mee and Yeaman 2019).

Among the set of genes showing no significant evidence of local adaptation were the tree freezing tolerance CBF transcription factors and CBF-regulon genes exhibiting GxE interactions. Among the GxE-CBF-regulon genes with cis-regulatory/nonsynonymous SNPs (51/53), only 20% contained high AFD SNPs (>0.60), which is similar to the proportion observed by E genes (Figure 2a). Furthermore, high AFD and/or LD GxE-CBF-regulon genes did not show significant enrichment along genetic-tradeoff QTL (Table S2). Finally, we did not find any significant evidence of local adaptation (i.e., in AFD and/or LD) along the three CBF transcription factors, supporting evidence that lower expression of these genes in warmer climates is under relaxed (Zhen and Ungerer 2008b; Zhen, et al. 2011) and not positive selection (Monroe, et al. 2016). It will be quite surprising if there is a unique genomic signature of adaptation underlying the three CBF genes, since high AFD and LD SNPs were enriched along the same genetic tradeoff QTL (GT QTL 4:2) but in a different region (Price, et al. 2020).

Interestingly, we find that GxE genes showing evidence of adaptation, show high expression and selective constraint/purifying selection—a pattern observed in other species (Wollenberg Valero, et al. 2014; Maddamsetti, et al. 2017; Boissot, et al. 2020). An interesting hypothesis that could be tested in future studies using Crispr-Cas9 technology (Cong, et al. 2013), is whether the effect size of candidate adaptive variation is positively correlated with levels of expression and selective constraint. This is partially supported by our study, since genes underlying the small number of high effect size QTL (Table 2) show higher mean expression than other genes. Unfortunately, we could not compare selective constraint between genes in Table 2 and the genome average given the very small sample of genes (n=9) with orthologs in *A. halleri*.

In conclusion, our study shows how genomic signatures of local adaptation, recent selection, and selective constraint, are linked to expression and fitness variation between Italy and Sweden ecotypes. Temperature is one of the environmental variables that may underlie local adaptation underlying Arabidopsis populations, therefore, examining more variables (e.g., precipitation (Postma, et al. 2016; Exposito-Alonso, et al. 2018; Monroe, et al. 2018)) may help us further understand the genetic architecture of local adaptation (Dittmar, et al. 2016) and the selection forces underlying it.

## Notes

### Competing Interest Statement

The authors have declared no competing interest.

