## Supplemental Figures and Tables for "Linking genomic signatures of selection to expression variation and direct evidence of local adaptation"

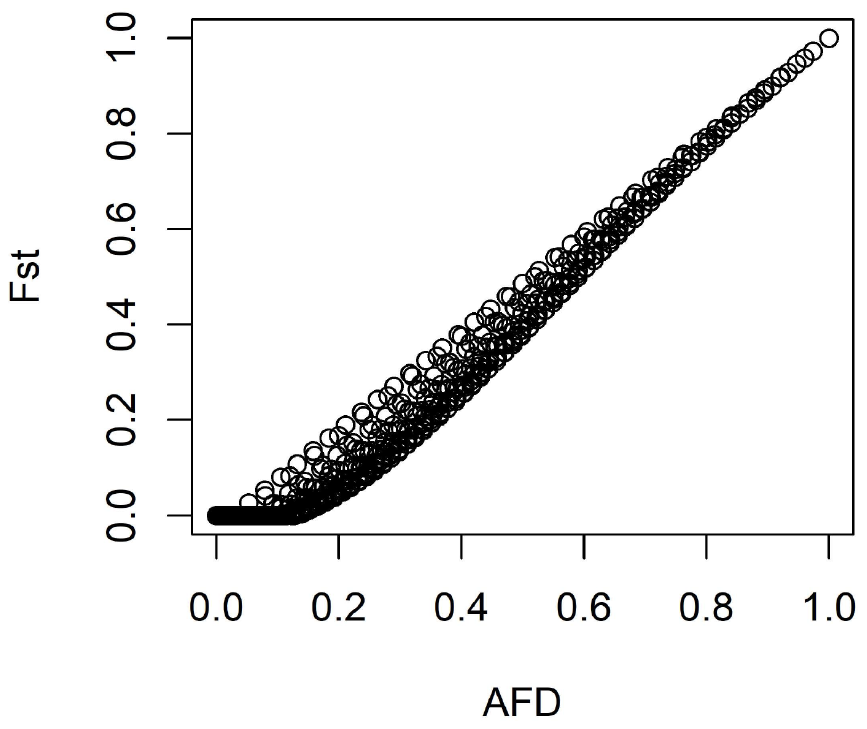

**Figure S1**. Comparing absolute allele frequency divergence between Italy and Sweden populations ($|f_{N.Sweden}-f_{S.Italy}|$ ) to an F_ST_  a two population, two allele, F_ST_ measure (Bhatia, et al. 2013). The estimated coefficient of determination (R^2^) was 0.96.

**Table S1**. Testing for enrichment of E and GxE genes exhibiting signatures of local adaptation and selection, within 100kb of 20 QTL peaks explaining fitness variation between Italy and Sweden populations (Ågren, et al. 2013). Compared to a genome-wide set of genes only GxE genes containing cis-regulatory/nonsynonymous SNPs with an AFD>0.60 & LD>0.32 showed a significant enrichment relative to the genome-wide proportion (*). This proportion of GxE genes was also significantly higher than the corresponding proportion of E genes (†). Comparison of proportions was done using a fisher’s one-tail test (p-val<0.05) implemented in R (“fisher.test”).

|  | NON-QTL | | | GT-QTL-PEAKS (100 kb) | | | E  (Odds ratio) | GxE  (Odds ratio) |
| --- | --- | --- | --- | --- | --- | --- | --- | --- |
|  | Genome-wide | E | GxE | Genome-wide | E | GxE |  |  |
| AFD>0.60 | 10867 | 925 | 152 | 558 | 55 | 7 | 1.16 | 0.90 |
| AFD>0.60 & LD>0.19 | 1487 | 136 | 18 | 141 | 18 | 4 | 1.40 | 2.34 |
| AFD>0.60 & LD>0.32 | 238 | 27 | 3 | 25 | 4 | 3 | 1.41 | 9.36*† |

**Table S2**. Testing for enrichment of CBF-regulon genes (Park, et al. 2018) exhibiting genomic signatures of local adaptation and selection; within six genetic-tradeoff QTL (Ågren, et al. 2013). Among three sets of genes: (i) the whole set of CBF-regulon genes (CBF); (ii) CBF-regulon genes showing main effect in environment (CBF-E); and (iii) CBF-regulon genes exhibiting genotype by environment interactions (CBF-GxE), none of them showed a significant enrichment withing GT QTL when compared to a genome-wide set of genes Comparison of proportions was done using a fisher’s one-tail test (p-val<0.05) implemented in R (“fisher.test”).

|  | NON-QTL | | | | GT-QTL | | | | ALL (OR) | E  (OR) | GxE  (OR) |
| --- | --- | --- | --- | --- | --- | --- | --- | --- | --- | --- | --- |
|  | Genome-wide | CBF | CBF-E | CBF-GxE | Genome-wide | CBF | CBF-E | CBF-GxE |  |  |  |
| AFD>0.60 | 9082 | 165 | 7 | 17 | 2343 | 38 | 2 | 3 | 0.89 | 1.11 | 0.68 |
| AFD>0.60 & LD>0.19 | 1209 | 21 | 1 | 2 | 419 | 7 | 0 | 0 | 0.96 | 0 | 0 |
| AFD>0.60 & LD>0.32 | 192 | 2 | 0 | 0 | 71 | 2 | 0 | 0 | 2.69 | 0 | 0 |

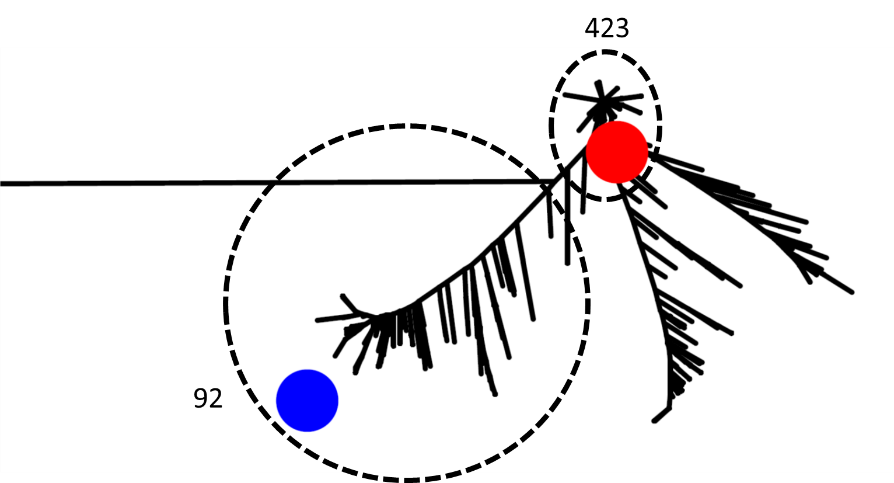

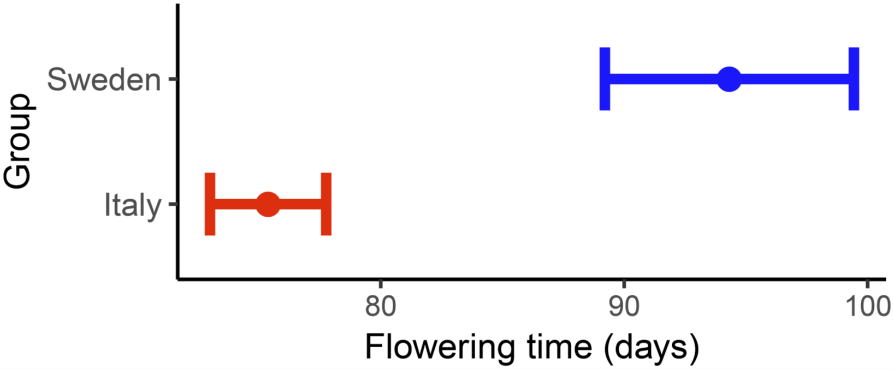

**Figure S2**. The rooted gene tree of COL7 using 875 *A. thaliana* Eurasian accessions and outgroups *Arabidopsis lyrata* and *Capsella rubella*. Eurasian accessions with a similar genotype as the Sweden parent (blue dot) that was used to identify fitness QTL in Ågren et al. (2013) show later flowering times than Eurasian accessions with a similar genotype as the Italy parent (red dot). The plot depicts the mean flowering time and 95% CI’s of 92 accessions with a similar genotype as the Sweden parent and 423 accessions with similar genotype as the Italy parent.

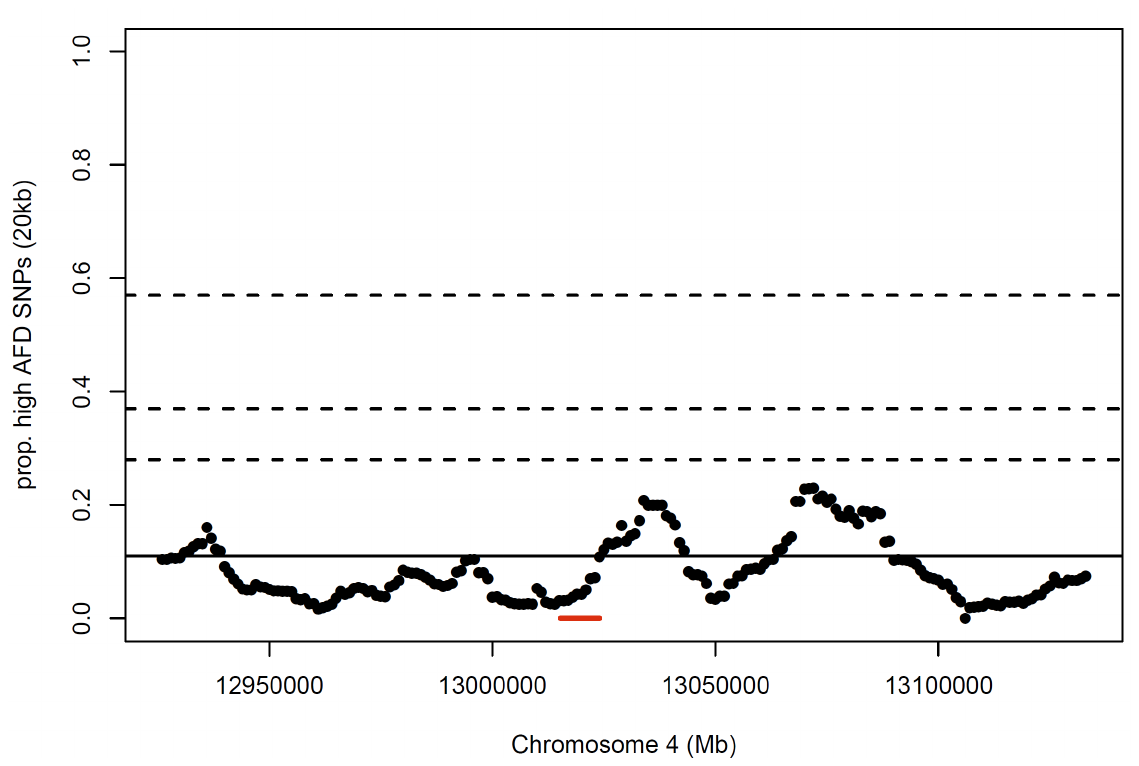

**Figure S3**. The genomic region spanning the three CBF transcription factors (CBF’s 1-3) and including candidate cis-regulatory regions (1kb distance from transcriptional start site) (shown in a red line), did not show a significant proportion of SNPs showing a high AFD (>0.60). Shown are the proportion of SNPs within a 20kb window that showed an AFD>0.60. The solid line indicates the genome-wide average, and the three dotted lines the 90^th^, 95^th^, and 99^th^ percentile of the genome-wide distribution of proportions.
